# Slow rhythmic activity from an interplay of voltage and extracellular concentration dynamics: a minimal biophysical mechanism for neuronal bursting

**DOI:** 10.1101/2023.12.12.571202

**Authors:** Mahraz Behbood, Jan-Hendrik Schleimer, Susanne Schreiber

## Abstract

Slow brain rhythms, for example during slow-wave sleep or pathological conditions like seizures and spreading depolarization, can be accompanied by synchronized oscillations in extracellular potassium concentration. Slow brain rhythms typically have longer periods than tonic action-potential firing. They are assumed to arise from network-level mechanisms, involving synaptic interactions and delays, or from intrinsically bursting neurons equipped with ion channels of slow dynamics. Here, we demonstrate that both mechanisms are not necessarily required and that slow rhythms can also be generated from an interplay of fast neuronal voltage dynamics and changes in extracellular ionic concentrations alone in any neuron with type Ⅰ excitability. The coupling of fast-spiking neuron dynamics and a slow extracellular potassium transient is regulated by the Na^+^/K^+^-ATPase. We use bifurcation analysis and the slow-fast method to reveal that this coupling suffices to generate a hysteresis loop organized around a bistable region that emerges from a saddle-node loop bifurcation – a common feature of type Ⅰ excitable neurons. Moreover, the Na^+^/K^+^-ATPase not only plays a key role in burst generation by shearing the bifurcation diagram but also modulates tonic spiking and depolarization block by its density and pump rate. These dynamics of bursting, tonic spiking and depolarization block, accompanied by the fluctuation of extracellular potassium, are likely to be relevant for pathological conditions. We suggest that these dynamics can result from any disturbance in extracellular potassium regulation, such as glial malfunction or hypoxia. The identification of a minimal mechanistic requirement for producing these dynamics adds to a better understanding of pathologies in brain rhythms may direct attention to alternative pharmacological targets for therapy.

**Author Summary:** The brain can produce slow rhythms, such as those observed during sleep or epilepsy. These rhythms are much slower than the neuronal electrical signals, and their origins are still under debate. Mechanisms discussed so far are based on the connection delays in neural networks or on neuronal ion channels with particularly slow kinetics. We show that neurons with specific spiking dynamics – allowing them to fire at arbitrarily low frequencies (type Ⅰ neurons) – can produce slow rhythmic patterns without requiring synaptic connectivity or special ion channels. In these cells, slow rhythmic activity arises from the interplay of slow changes in extracellular potassium concentration and the cell’s voltage dynamics, mediated by the Na+/K+-ATPase pump. The latter, found in all neurons, regulates the concentrations of sodium and potassium ions across the cell membrane. The core mechanism is not idiosyncratic, rather mathematical analysis shows under which conditions slow rhythmic activity can arise generically from the pump-based coupling in a broad class of neurons. We demonstrate that the pump is relevant for the creation of different firing patterns, which can be associated with various diseases. A better understanding these complex dynamics is important for the development of more effective treatments for concentration-dependent pathologies.

## Introduction

Rhythms on diverse timescales are a hallmark of our brain’s activity. Such oscillatory activity is associated with physiological states, ranging from normal breathing to the consolidation of memories, or pathological situations such as epileptic seizures and spreading depolarization (SD)[1–5]. Some rhythms are orders of magnitude slower than single-neuron spiking. For these, different mechanistic origins of their slow dynamics have been proposed. First, a neuron’s embedding in the network can produce bursts via reverberating activity, with synaptic delays serving as an important parameter that determines the period of the resulting oscillations [6]. Second, one or more cells with intrinsically bursting dynamics can act as pacemakers [7] and entrain larger networks. Combinations of both of the above mechanisms, network effects and intrinsic bursting, have also been reported [8].

Brain regions that exhibit large-scale oscillatory behaviour often contain significant numbers of single-neuron bursters [9–11], fostering network entrainment via pacemaker neurons [3]. Theoretical and experimental research suggests that the neuron’s ability to generate bursts of activity intrinsically arises from the specific ion channels expression, such as hyperpolarization-activated inward transient potassium channels, or calcium-dependent channels [3,12–14]. The kinetics of these channels form the basis of the slow dynamics.

Ion channels with specialised kinetics, however, are not the only single-cell mechanism that can generate rhythms on a slow timescale. In this study, we argue that, in principle, any type Ⅰ nerve cell can be switched into an intrinsic burster even in the absence of channels with slow dynamics via the interplay of cellular spiking and extracellular concentration dynamics. For the proposed mechanism two facts are combined : (1) All neuron models which can produce arbitrarily slow firing rates (type Ⅰ excitable neurons [15,16]) can be tuned into a bistable regime (where a stable fixed point and a limit-cycle co-exist) [17–19]. Alternating between the two stable stages in the bistable regime is the basis of the deterministic bursting mechanism. (2) The oscillation between two distinct dynamics within the bistable region is orchestrated by the interplay between the neuron’s fast-spiking mechanism and the comparatively slow evolution of ionic concentrations. This study concentrates on extracellular potassium dynamics as the slow mechanism contributing to the emergence of bursting behaviour. Moreover, the interaction between extracellular potassium concentration and the spiking mechanism is regulated by feedback from the Na^+^/K^+^-ATPase pump which is responsible for the homeostasis of sodium and potassium concentration in the neurons [20]. The slow rhythm produced by this mechanism is not only expressed at the level of neuronal spiking but also ionic concentrations.

Experimental findings indicate a harmonious oscillation between extracellular potassium concentration and the rapid spiking changes during bursting [21,22]. Moreover, specifically, potassium concentration-based bursting has been noted in complex models [12,23]. Here, the generality of such a mechanism is analysed, based on a parsimonious model comprised of only the most essential ingredients. These encompass a variable extracellular potassium concentration in an extracellular environment of fixed size impacted by the activity of the Na^+^/K^+^-ATPase as well as classical Na- and K-channel-based spiking of the neuron. We show that for such a rhythmic slow bursting, no special channels or rhythmic input from a network is required.

In the following, a model exemplifying the proposed generic mechanism of slow bursting that arises from the interplay of fast spiking and slow extracellular potassium concentration dynamics is introduced. Next, the mechanism underlying the slow deterministic bursting is analysed using the dissection of slow and fast dynamics (here also termed the slow-fast method) [24]. Based on numerical continuation, the bifurcations in the fast subsystem are identified and the existence of a slow hysteresis loop mechanism is demonstrated. Finally, the conditions for which the model can be tuned into the slow bursting regime or other dynamics such as depolarization block and tonic spiking are discussed. Taken together, the proposed mechanism highlights the crucial role of extracellular potassium dynamics and Na^+^/K^+^-ATPase in the emergence of slow rhythms at the single neuron level and beyond.

## Results

### Modelling neurons in an extracellular environment

In order to study the emergence of slow rhythmic activity via the interactions of fast spiking and slow concentration dynamics (i.e., fast and slow subsystems, respectively. see next sections), a neuron model equipped with only basic fast spike-generating ion channels is placed in an external environment of fixed volume that contains extracellular potassium ions, whose concentration, [K^+^]_out_, can change. The interplay between concentration dynamics and neuronal voltage dynamics is modulated by the activity of the electrogenic Na^+^/K^+^-ATPase pump and the influence of ion flow across the neuronal membrane on the concentration-dependent reversal potentials, see Fig 1. It will be shown in the subsequent section that this parsimonious setup suffices to create slow rhythms and of what nature these rhythms are.

**Fig 1.**
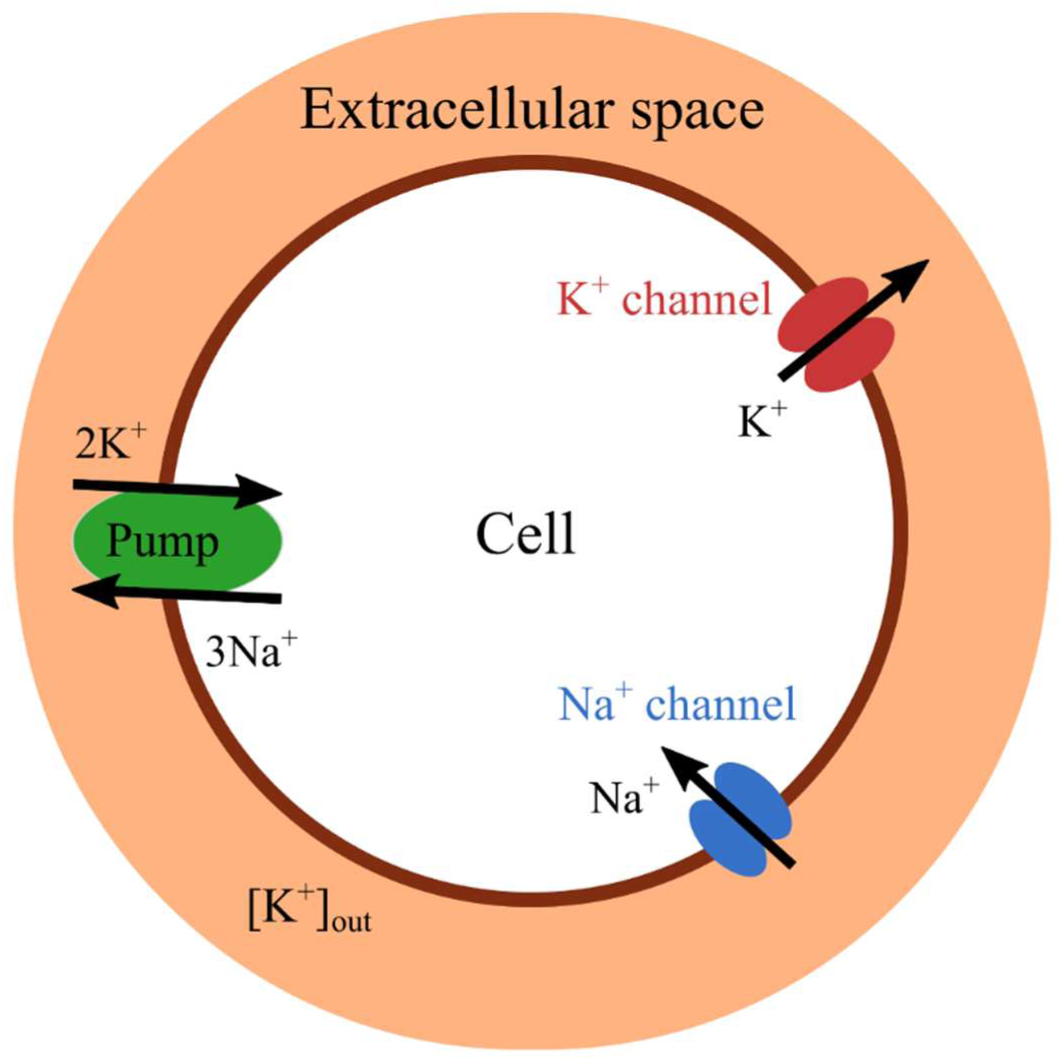
Schematic drawing of the neuron model in the extracellular environment. The concentration of extracellular potassium changes with the activity of voltage-gated potassium channels (red) and the Na^+^/K^+^-ATPase pump (green). The model equations governing the spike-generating inward sodium (blue) and outward potassium (red) currents correspond to those in the original Wang-Buzsáki model, retaining their fast dynamics. A slower timescale arises from the dynamics of the extracellular potassium concentration.

The neuron model, inspired by the Wang-Buzsáki model, comprises four currents: The depolarising Na^+^-current and repolarising K^+^-current are part of the spike-generating mechanism. A leak current balances the membrane potential in the rest state. For completing our model, we add a fourth current as a hyperpolarising Na^+^/K^+^-ATPase pump current (I_p_). This pump extrudes three Na^+^ ions out of the cell while two K^+^ cations are pumped in. The pump maintains the ionic gradient of the cell, which drives spiking activity. The activity of the pump is regulated by the extracellular potassium concentration, with higher [K^+^]_out_ resulting in increased pump strain (Eq 7). However, the pump’s activity is limited by its maximum current capacity (I_max_), which is influenced by both the electrophysiology characteristic of the pump and its density on the neuron’s membrane. For the mathematical details, see Methods.

With each action potential, K^+^ ions flow out of the cell through the voltage-dependent K^+^ channels and accumulate in the extracellular space. The elevated extracellular potassium concentration ([K^+^]_out_), in turn, not only affects the pump current but also influences the neuron’s excitability state via its effect on the reversal potential. Therefore, the dynamics undergoes a sequence of transitions, which form the basis for the rhythmic bursting behaviour and are explained and analysed in more detail in the following sections.

### Pump- and extracellular potassium-mediated slow bursting

In this section, we first provide a brief illustrative description of the generic burst mechanism, leaving a detailed analysis of the conditions for its existence to the following sections.

An example trace of the slow bursting dynamics that the neuron can produce in a dedicated extracellular volume is depicted in Fig 2A. Each burst cycle contains two alternating states: quiescence and spiking. The rhythmicity of the voltage shows an inter-burst period of about one second. This period is ≍30 times longer than the dynamics of one complete action potential in the spiking phase. The number of spikes per burst is fixed (i.e., bursting is deterministic).

**Fig 2.**
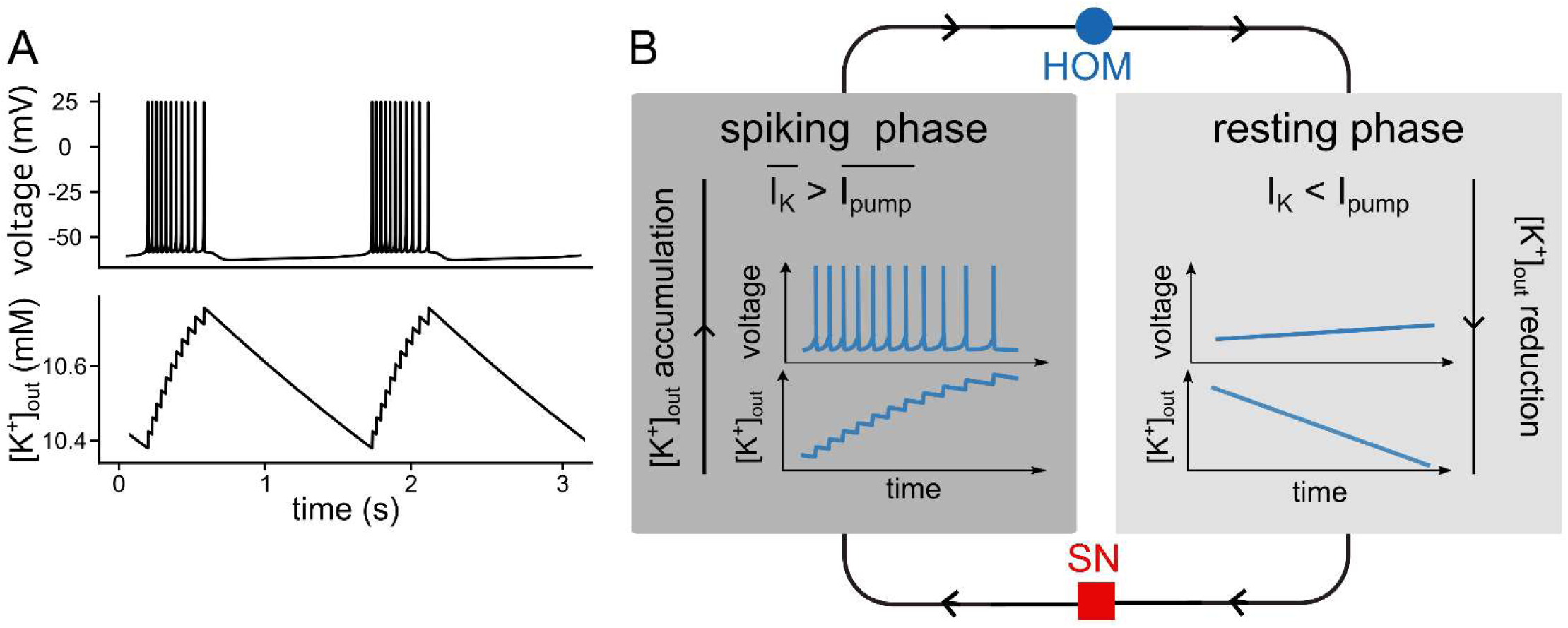
Bursting dynamics overview. (**A**) Each bursting cycle consists of two phases: a spiking phase and a quiescent phase, visible in the voltage trace. The concentration waxes with each spike and wanes during the quiescent phase. (**B**) The spiking phase commences at a saddle-node (SN) bifurcation of the fast subsystem (red square). During the spiking phase, the average outward potassium flux through potassium channels surpasses the average inward potassium flux through the Na^+^/K^+^-ATPase leading to an accumulation of extracellular potassium. This phase terminates at a homoclinic (HOM) bifurcation of the fast subsystem (blue circle). From this bifurcation, the quiescent phase initiates, accompanied by a continuous decrease in extracellular potassium concentration due to the Na^+^/K^+^-ATPase current superseding the potassium current through the voltage-gated channels. This eventually results in a new spiking phase. I_app_=0.5 μA/cm^2^ and I_max_=1 μA/cm^2^.

The basic cycle undergone during a burst is summarised in Fig 2B. The burst relies on the bistable nature of the neuron’s dynamics between spiking and quiescence. During the spiking period, the outward flow of potassium ions is considerable. The inter-spike interval does not provide sufficient time for the Na^+^/K^+^-ATPase to pump back all K^+^ ions that left the cell during the spike. Consequently, K^+^ ions accumulate in the extracellular space, resulting in a saw-tooth-like build-up of [K^+^]_out_ (Fig 2B, spiking phase). Furthermore, during the spiking phase of each burst, the inter-spike interval progressively prolongs as a result of K^+^ accumulation in the extracellular space. Eventually, the rise of [K^+^]_out_ reaches a level where spiking is not possible any more (the quiescent phase). Spiking is terminated by a homoclinic bifurcation (HOM) in the fast subsystem, indicated by the blue circle in Fig 2B. In the quiescent phase, the pump activity now suffices to reduce extracellular potassium, as the outward current via the potassium channels is significantly reduced in the absence of spike generation. Ultimately, the decrease of [K^+^]_out_, during quiescence results in another bifurcation of neuronal dynamics, corresponding to saddle-node bifurcation (SN) of the fast subsystem, see red square in Fig 2B. In passing this bifurcation, neuronal dynamics are back in the spiking region and the cycle starts all over again.

In summary, the spiking dynamics of the neuron impacts [K^+^]_out_, the latter of which, in turn, influences the K^+^ reversal potential and hence the neuron’s voltage dynamics. In the next sections, it will be shown that the identified mechanism is, indeed, a *hysteresis loop* where one branch of the bistable system corresponds to the spiking state and the other branches constitutes the quiescent state. The properties of the hysteresis loop are then further analysed in the slow-fast analysis section. The remainder of the article focuses on establishing how generic this mechanism is.

### Slow-fast analysis

Neuronal burster consists of an oscillation governed by two time scales: one corresponding to the fast spiking dynamics and one corresponding to the slower dynamics related to concentration changes. Typically, these two different time scales result from either the presence of fast and slow currents [24], stochastic escape [25], external drive, synaptic delays [6], or as here, the slow variation of ions. All these cases benefit from a slow-fast analysis [24].

In the mechanism for the generation of slow rhythms proposed here, the fast subsystem is governed by the millisecond kinetics of the ion channels that shape the neuron’s action potentials. In contrast, the slow subsystem is governed by the timescale of slow changes in [K^+^]_out_. The bursting rhythmicity of the system is thus regulated by an interplay between these two subsystems, which crucially depends on the activity of the Na^+^/K^+^-ATPase.

In slow-fast analysis, it is assumed that the timescale difference between the two subsystems is sufficiently large for them to be investigated separately. We will first subject the fast subsystem to bifurcation analysis while assuming that the slow subsystem undergoes only negligible variations on the timescales of the fast subsystem. Hence, [K^+^]_out_ can be treated as constant and the bifurcation parameter in this scenario. Subsequently, we will explore the dynamics of the slow subsystem assuming time-averaged values of the fast subsystem dynamics. More details on the method are to be found in the Methods section.

#### Fast subsystem bifurcation analysis

For analysis of the fast, spike-generating subsystem, extracellular potassium concentration is approximated as a constant parameter. The bifurcation diagram in Fig 3 summarises the transitions in dynamics induced by changes in [K^+^]_out_.

**Fig 3.**
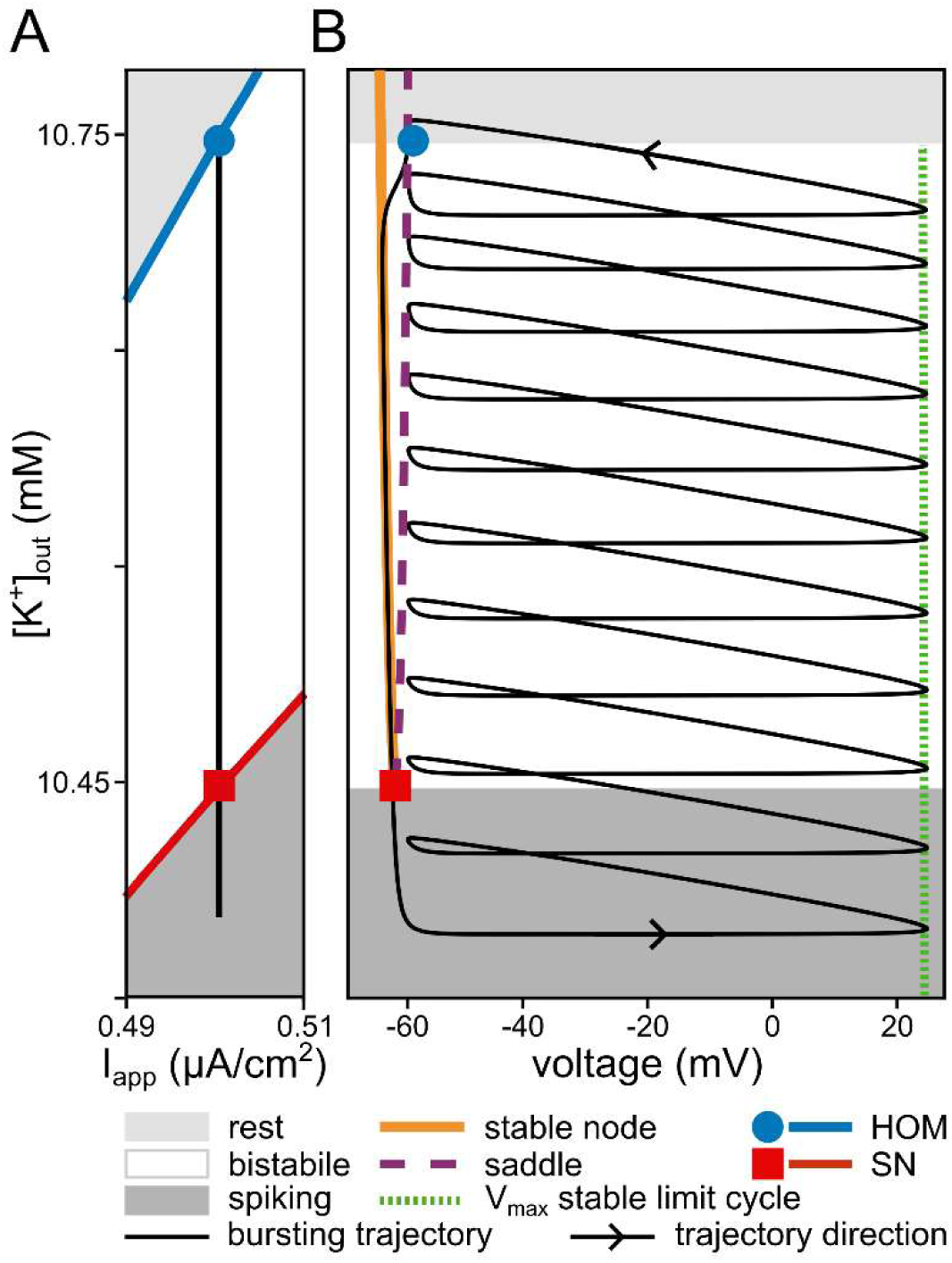
Bifurcations of the fast subsystem and bursting dynamics. The bistability of the fast subsystem is a requirement for the emergence of bursting dynamics in our system. (**A**) Two-parameter (extracellular potassium, [K^+^]_out_, and applied current, I_app_) bifurcation diagram of the fast subsystem. The black vertical line indicates the range of [K^+^]_out_ values that the complete system traverses during the burst cycle shown in Fig2A. The burst trajectory passes through the bistable region, bounded by the homoclinic (HOM) bifurcation at higher [K^+^]_out_ and the saddle-node (SN) bifurcation at lower [K^+^]_out_. During the spiking phase of the burst, [K^+^]_out_ builds up, whereas it depletes during the resting part. The black line extends into the tonic spiking domain of the fast subsystem due to the time spent near the ghost of the SN bifurcation. (**B**) Phase portrait of the complete burst (in voltage and [K^+^]_out_) onto the one parameter ([K^+^]_out_) bifurcation diagram of the fast subsystem. The black solid line demonstrates the same bursting dynamics as in Fig2A.

Fig 3A shows a two-parameter bifurcation diagram of the fast subsystem spanned by the parameters extracellular potassium ([K^+^]_out_) and applied current (I_app_) in the vicinity of the region used for the simulated traces in Fig 2A. Overlaid on the bifurcation diagram of Fig 3A are the dynamic changes of the complete system during a burst cycle with the parameters of Fig 2A shown as the black vertical line. If [K^+^]_out_ is low, the only stable state for the fast subsystem is a stable limit cycle (the dark grey area in both panels of Fig 3). In this region, the neuron spikes tonically. If [K^+^]_out_ is increased, the dynamics of the fast subsystem traverse the saddle-node (SN) bifurcation (the red line in Fig 3A). From the SN bifurcation, a saddle and a stable node emerge. When [K^+^]_out_ exceeds the SN bifurcation value, the neuron enters the bistable region (the white region in both panels of Fig 3). In the bistable region, the fast subsystem of the neuron model has two stable states, a stable limit cycle and a stable fixed point, separated by a separatrix originating from the stable manifold of the saddle. By raising [K^+^]_out_ even more, the blue line of a homoclinic orbit bifurcation (HOM) is crossed, at which the limit cycle of the action potential collides with the saddle and is annihilated. If [K^+^]_out_ passes this line, the fast system dynamics end up in the monostable zone, because at the HOM bifurcation, the stable limit cycle disappears. Hence, in the light grey part of Fig 3A, the neuron is in a quiescent state.

To better understand the bursting dynamics, the voltage trajectory of one burst cycle in the complete system (black solid line) is superimposed on the codimension-one ([K^+^]_out_) bifurcation diagram of the corresponding fast subsystem (coloured lines) in Fig 3B. The black trajectory depicts the burst dynamics presented in Fig 2A, however, instead of time the corresponding levels of [K^+^]_out_ are shown. The resulting phase portrait encompasses the complete system dynamics of the burst: In the dark grey region of Fig 3B the fast subsystem is in the tonic spiking mode. Here, the complete system (black trace) is going through a spiral path. Each rotation corresponds to one action potential. With each action potential, more K^+^ accumulates in the extracellular space (see also Fig 2). By increasing [K^+^]_out_, the system enters the fast subsystem’s bistable region (white area) and continues on the spiking branch. Note that if the neuron model was subjected to strong perturbations, jumps between the two attractors, a stable fixed point and a stable limit cycle, would occur. The effective increase in [K^+^]_out_ per spike diminishes as spiking progresses because the pump becomes more and more active due to the increase of extracellular potassium (Eq 7). Furthermore, the inter-spike interval increases because of the changes in the spiking dynamics caused by changes in the potassium reversal potential due to the accumulation of extracellular potassium.

Spiking continues until, eventually, [K^+^]_out_ reaches the HOM bifurcation (blue circle) and the stable limit cycle of the fast subsystem is annihilated. Here, the system enters the quiescent phase of the fast subsystem (light grey zone). In this phase, the only stable state is the rest state. Hence, the system trajectory leaves the spiking spiral and converges to the stable fixed point (orange line). In the rest state, [K^+^]_out_ is efficiently pumped back into the cell by the Na^+^/K^+^-ATPase (see also Fig 2). The pump activity increases the resting state voltage until an SN bifurcation is reached (red square). The saddle and the stable node collide and annihilate at this point. Thereby, the resting state loses stability and the system converges back onto the tonic spiking dynamics (dark grey), where the next burst cycle is initiated; for a zoom out of the bifurcation diagram in Fig 3B see S 1 Fig).

Notable in both panels of Fig 3, there is a gap between the SN bifurcation and the onset of the first spike in a burst cycle. In this region, the voltage dynamics cannot, after passing the SN bifurcation, immediately jump to the spiking branch due to the proximity to the ghost of the SN bifurcation (i.e., the dynamics is slowed down in this region). Thus, the decrease of the [K^+^]_out_ does not immediately stopped when the system reaches the SN bifurcation. [K^+^]_out_ is further reduced when moving away from the SN ghost until it can finally enter the spiking phase.

#### Slow subsystem analysis

The previous section analysed the spiking behaviour with the slow variable, specifically, the extracellular potassium concentration ([K^+^]_out_) treated as a parameter. The fast subsystem bifurcations that initiate and terminate spiking were identified. In this section, the dynamics of the slow subsystem is explored. The time scale of the slow subsystem is determined by the effect of spiking on [K^+^]_out_ in relation to the balancing effect of the Na^+^/K^+^-ATPase. Note that the effective ATPase pump rate depends on both the extracellular potassium concentration as well as the density of ATPase proteins (indirectly defined by I_max_, see the Eq 7). Relevant quantities for the slow subsystem are thus the amount of K^+^ leaving the cell per spike and the effective pump rates during the spiking and quiescent phases.

The slow concentration dynamics can be isolated by applying the method of averaging to the fast dynamics. It exploits the effect that the slow subsystem cannot track the rapid changes in the fast subsystem; effectively, it only senses the average impact of one complete spike. To calculate this average impact on the change of [K^+^]_out_, the pump current and the current carried by the fast spiking potassium channels are integrated over one spike period. The averaged quantity of the fast subsystem currents determines the time derivative of the slow subsystem and, consequently, its dynamics (see Eq 14). Note that when the fast subsystem is quiescent (i.e., no spiking), averaged and non-averaged fast subsystem dynamics are equal. With this averaging method, we can describe the dynamics of the slow subsystem as a reduced model that can capture the complete system dynamics. See Methods for more details on the averaging method and [24] for an introduction.

Fig 4 shows the phase portrait of the reduced slow subsystem, in terms of [K^+^]_out_ and its temporal derivative, for the specific parameters used in Fig 2. In analogy to Fig 3, in Fig 4 areas of tonic spiking (dark grey), bistability (white), and quiescence (light grey) of the fast subsystem are marked. The HOM and SN bifurcations of the fast subsystem mark the borders of these zones, (blue and red horizontal lines). Trajectories of the reduced slow subsystem corresponding to spiking and quiescence of the fast subsystem are indicated by the labels ‘spiking’ and ‘rest’, respectively.

**Fig 4.**
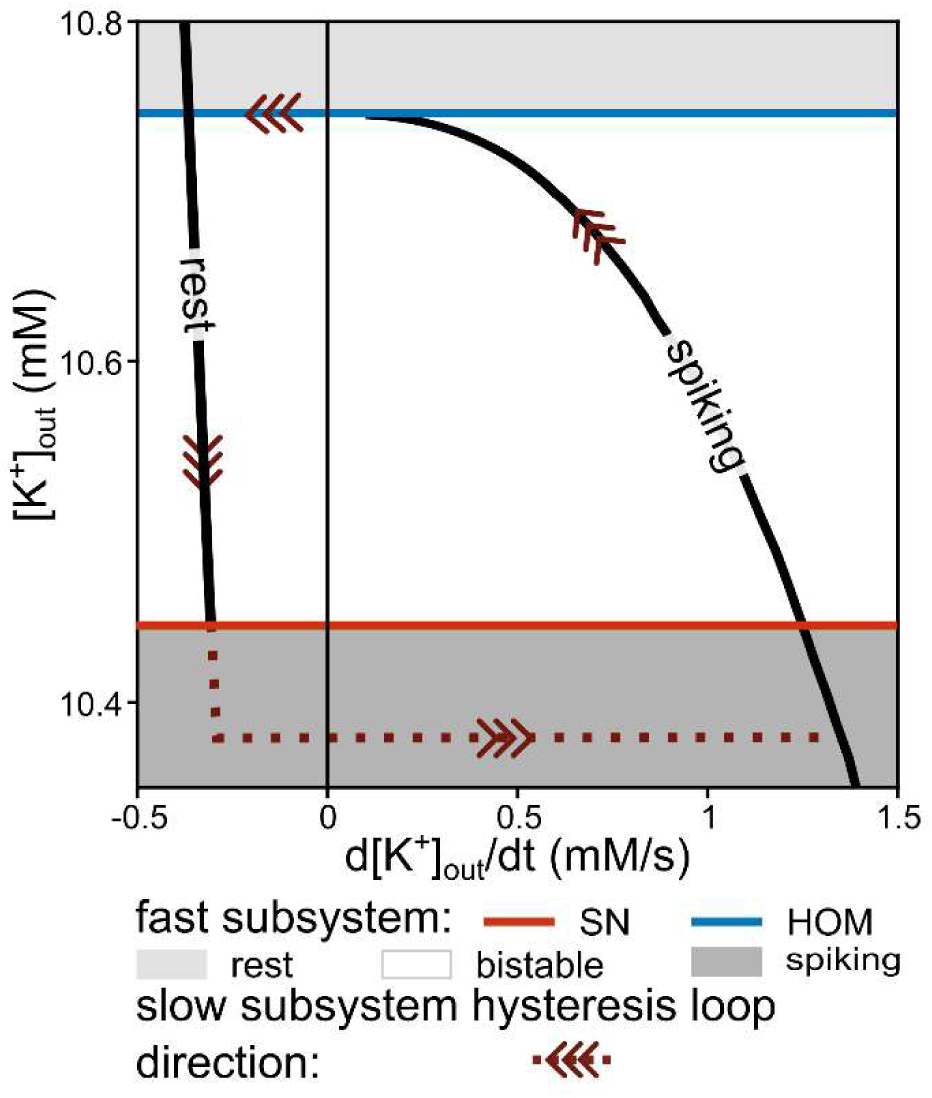
Hysteresis loop of the slow subsystem overlaid with the bifurcations of the fast subsystem. The extracellular potassium ([K^+^]_out_) dynamics of the reduced, slow subsystem entails a hysteresis loop oscillator. The slow oscillation organises around the bistable region of the fast subsystem, as also shown in Fig 3. The slow subsystem trajectory for the resting branch is calculated by inserting steady state fast variables as a function of the slow concentration into the [K^+^]_out_ time derivative (see Methods, Eq. 9), while the spiking branch requires averaging (see Methods, Solow-fast method, Eq. 11). The red and blue lines represent the location of the saddle-node (SN) and homoclinic (HOM) bifurcations of the fast subsystem according to ([K^+^]_out_), respectively. Note that, after going through the SN bifurcation, the system remains near the SN ghost for a while before spiking is resumed (brown dotted line). Parameters as in Fig 2A.

Fig 4 reveals the hysteresis nature of the slow subsystem (reduced system), which revolves around the bistability of the fast subsystem. The bistability of the fast subsystem serves as the prerequisite for the existence of the hysteresis loop in the reduced slow subsystem. For clarification of the mechanism, we follow the [K^+^]_out_ hysteresis loop in Fig 4 through the same sequence of bifurcations as in Fig 3B. If our fast subsystem is spiking (equivalent to the spiral path in Fig 3B), this corresponds to the spiking branch of the hysteresis loop in Fig 4 and the time derivative of [K^+^]_out_ is positive. This means that in each cycle of an action potential, the mean K^+^ flux into the extracellular space is larger than the mean of the K^+^ flux via the pump, which reabsorbs potassium ions into the neuron. In other words, extracellular K^+^ accumulates continuously in extracellular space. However, while the slow subsystem slides on this branch from left to right through the bistable region, the potassium increment reduces with every additional spike. With the parameters used here, the spiking branch of the slow subsystem terminates at the HOM bifurcation before the derivative of [K^+^]_out_ turns negative. This observation depends on the system’s parameters, as it is further explored below. After the HOM bifurcation, the slow subsystem falls back to the stable resting branch.

In the quiescent state of the fast subsystem (light grey region), the K^+^ flux via the ATPase dominates, as there is no spiking and, consequently, no major flux of potassium out of the neuron. Therefore, the time derivative of [K^+^]_out_ is negative (Fig 4, rest branch). [K^+^]_out_ decreases and potassium is pumped back into the neuron. The slow subsystem follows the quiescence branch from higher to lower [K^+^]_out_ through the bistable zone (equivalent to sliding on the orange line in Fig 3B). At the SN bifurcation (red line), the stable node disappears from the fast subsystem’s dynamics. As a consequence, the silent branch comes to an end at a lower [K^+^]_out_. Subsequently, the reduced slow subsystem undergoes a transition to the spiking branch after passing through the ghost of the SN bifurcation. This completes the hysteresis loop of the reduced slow subsystem.

In summary, the bursting process is enabled by the bistability of the system and corresponds to a hysteresis loop. The bistability allows the system to alternate between quiescence (where the activity of the ATPase suffices to extrude potassium from the extracellular space) and spiking (marked by accumulation of potassium in the extracellular space).

### Generic occurrence of the burst mechanism

The previous sections described how, for specific parameter settings, type Ⅰ conductance-based neuron models can exhibit slow bursting when the dynamics include the activity-dependent concentration changes in an extracellular space of fixed volume. The following section investigates the robustness and prevalence of the described burst mechanism and its dependence on model parameters.

#### The pump’s essential contribution to the bursting mechanism

As the mechanism of bursting relies on a region of bistability, one may note that the generic unfolding of a codimension-two saddle-node loop (SNL) bifurcation from the neuron model without a pump includes a region of bistability (Fig 5A) [26]. For the burst mechanism to work, however, the fast dynamics along the [K^+^]_out_ direction needs to be bounded by HOM and SN branches for higher and lower values of [K^+^]_out_, respectively (compare Fig 3A). This is possible, if the bifurcation diagram is shear transformed such that the HOM and SN branches bend to the right. Interestingly, such shearing occurs as a result of adding a K^+^-dependent pump (Eq 7) to the neuron model with an SNL bifurcation (see Fig 5B). The pump current acts as an applied current that depends on [K^+^]_out_; in our model it is a sigmoidal function with half activation at 11 mM. The pump induces a shear transformation of the canonical unfolding of the SNL bifurcation (Fig 5B). The larger the maximum pump current (I_max_), the larger the shear. The Na^+^/K^+^-ATPase density in the membrane (directly related to I_max_), hence is a crucial parameter for the emergence of bursting by the minimalistic mechanism proposed here.

**Fig 5.**
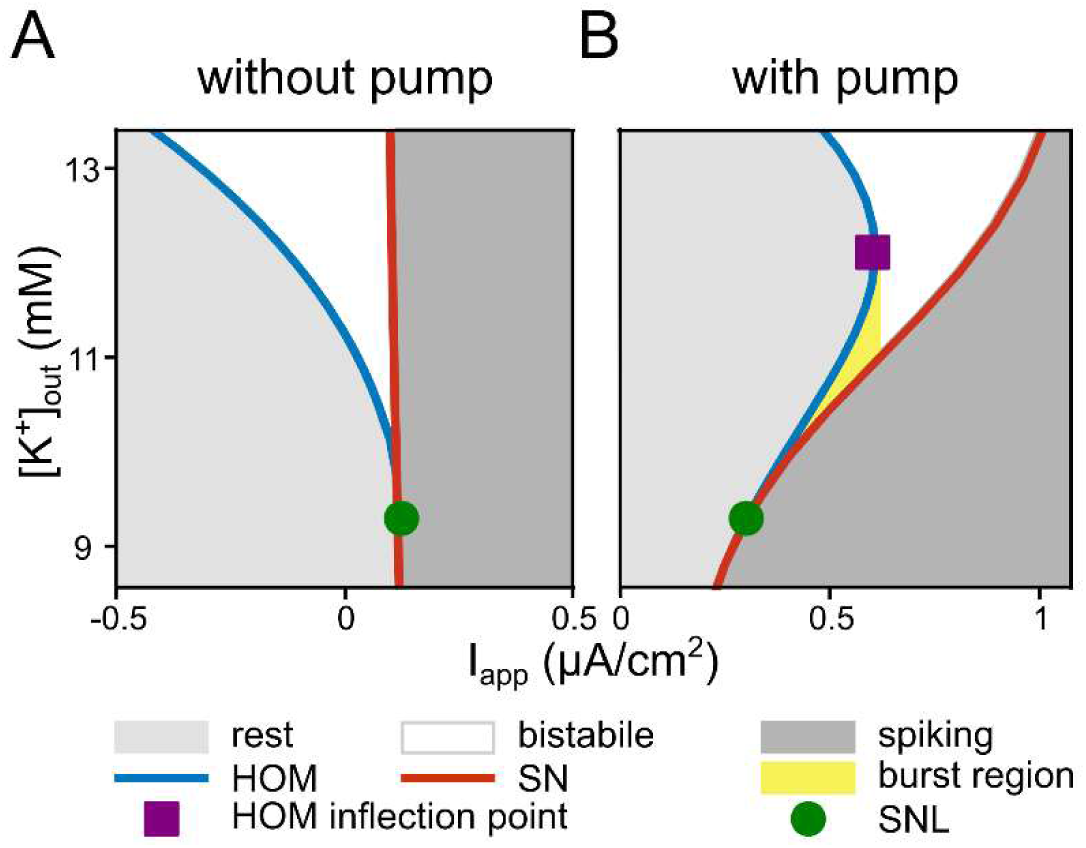
Shear transformation of the saddle-node loop (SNL) unfolding induced by the Na^+^/K^+^-pump. Including an Na^+^/K^+^-ATPase into the neuron model affects the fast subsystem bifurcation diagram and thereby enables bursting dynamics. (**A**) Two-dimensional bifurcation diagram of the fast subsystem without Na^+^/K^+^-ATPase. The generic unfolding of the SNL bifurcation (green circle) includes a bistable area between the homoclinic (HOM) and saddle-node (SN) bifurcation lines. (**B**) Same bifurcation diagram after adding Na^+^/K^+^-ATPase current to the model, which shears the area of bistability. This opens up the possibility for a [K^+^]_out_ hysteresis loops (yellow region) for values of I_app_, between SNL and HOM inflection point (purple square). I_max_=1 μA/cm^2^.

The region of burst occurrence in the fast subsystem (yellow area in Fig 5B) is bounded by the SNL point (green circle) and the inflection on the HOM branch (purple square). For I_app_ beyond the inflection point, spiking continues until [K^+^]_out_ reaches a fold of limit cycles (FLC), which is analysed in the next section.

In summary, the burst mechanism relies on two essential ingredients from the neuron’s fast subsystem, which enable the hysteresis loop:

i. For low [K^+^]_out_, the onset of spiking in the fast subsystem occurs via a saddle-node on invariant cycle (SNIC) bifurcation (typically present in neuron with class I excitability). This implies that the neuron model also undergoes an SNL bifurcation; at higher [K^+^]_out_ [27,28], opening up the bistable region between a homoclinic and a SNIC branch (Fig 5A).
ii. The electrogenic Na^+^/K^+^-ATPase counteracts the K^+^ efflux and mediates the coupling between [K^+^]_out_ and neuronal voltage dynamics. The pump’s activity shears the bistable region such that it can be traversed by changes in [K^+^]_out_ while being bounded by a fixed-point region (quiescence) from above and a limit-cycle region (tonic spiking) from below.

#### Dynamics in the proximity of the bursting region

To comprehensively understand the model’s dynamics, we need to take a detailed look at the dynamics occurring in the vicinity of the bursting in the full system to shed light on factors that constrain bursts, the dependence on parameters, and the robustness of the described mechanism.

Zooming further out of the K_out_-I_app_ plane depicted in Fig 3A reveals additional structures in which the region of bistability of the fast subsystem is embedded (Fig 6A). The vertical trajectories overlaid on the fast subsystem bifurcation show the direction of the slow subsystem (i.e., complete system) dynamics. According to Fig 6A, variation of the applied current resulted in different dynamics of the complete system. The slow bursting is possible in the range of I_app_ between SNL (green circle) and the HOM inflection point (purple square), as is the case in the yellow area in Fig 5B. The specific bursting dynamics of Fig 2A is found in the parameter region D in Fig 6A.

**Fig 6.**
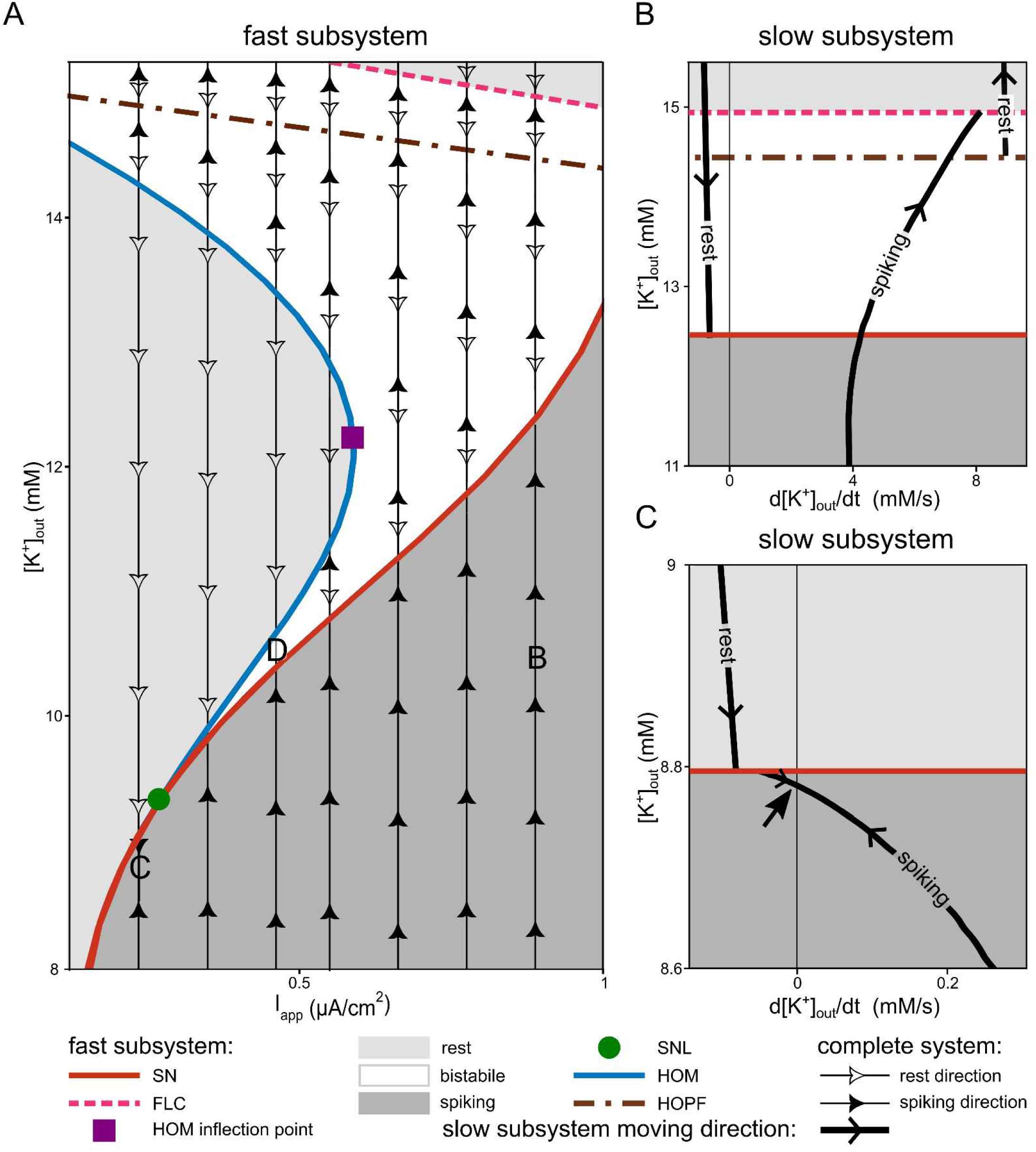
Bifurcations of the fast subsystem with an overlaid phase portrait around the bursting region. The bifurcations of the fast subsystem together with the flow of dynamics is shown in the vicinity of bursting area. (**A**) Zoom-out of the two-parameter bifurcation diagram of the fast subsystem depicted in Fig 3A. Depending on the applied current (I_app_) and extracellular potassium concentration ([K^+^]_out_), the fast subsystem exhibits mono-, bi-, or even multistability. The resting region in the top right corner, above the fold of limit cycles (FLC) is characterised by bistability between two fixed points. The region between the Hopf and FLC bifurcations is multistable between two fixed points and one limit cycle. The bistable region enclosed between the homoclinic (HOM), saddle-node (SN), and Hopf bifurcations enables bursting. Here, the bistability occurs between one limit cycle and one fixed point. The burst is possible for values of I_app_ between the ones corresponding to the saddle-node-loop (SNL) and HOM inflection point. The area marked as D highlights where the bursting shown in Fig 2A occurs. The vertical arrows indicate the flow field direction of the complete system. For the higher [K^+^]_out_ than the Hopf line, the one branch of the complete system flow field direction which is going to higher [K^+^]_out_ and indicating the depolarization block branch is not shown (see panel B, rest branch with positive time derivative of [K^+^]_out_). Two specific cases are selected from the region of depolarization block (B) and tonic spiking (C) of the complete system dynamics for further study. The reduced slow subsystem overlaid onto the bifurcation map of fast subsystem for these two points are depicted in panels B and C. (**B**) Reduced slow subsystem dynamics overlaid on the bifurcation of the fast subsystem in the case marked as B in panel A (I_app_=0.9 μA/cm^2^). This case corresponds to a transition to the depolarization block in the complete system. The arrangement of the resting and spiking branches indicates that, regardless of where the system starts on this diagram, it will eventually reach the resting branch with the higher time derivative of [K^+^]_out_ and follow that line up to the depolarization block. (**C**) Reduced slow subsystem dynamics overlaying on the bifurcation map of the fast subsystem in the case marked as C in panel A (I_app_= 0.25 μA/cm^2^). This case corresponds to tonic firing in the complete system. The black straight vertical line represents d[K^+^]_out_/dt =0, and the arrow indicates the spiking branch of the slow subsystem crossing this line and having a stable fixed point. Accordingly, the complete system remains tonically firing. I_max_ =1μA/cm^2^.

At I_app_ below the SNL bifurcation in Fig 6A, tonic spiking is possible. An example from the tonic spiking area is indicated by the letter C in Fig 6A. The bistable region, which is a requirement for the bursting mechanism under investigation, is absent in the fast subsystem bifurcation for this case. As shown in Fig 6A, in the vicinity of the position marked C, two dynamical regimes for the fast subsystem exist: tonic spiking and quiescence. Why has the complete system a stable state at tonic spiking? The answer can be read off Fig 6C, which depicts the time derivative of the reduced slow subsystem. The spiking branch of the reduced slow subsystem crosses zero with a negative slope (see arrow in Fig 7C); the reduced system has a stable fixed point. This fixed point is in the region where the only attractor in the fast subsystem is a stable limit cycle. The silent branch in Fig 6C is lower than zero. Consequently, if we start on this branch, the reduced slow subsystem eventually jumps to the spiking branch and reaches the stable fixed point. Thus, the complete system in the region around C ends up in the spiking regime.

**Fig 7.**
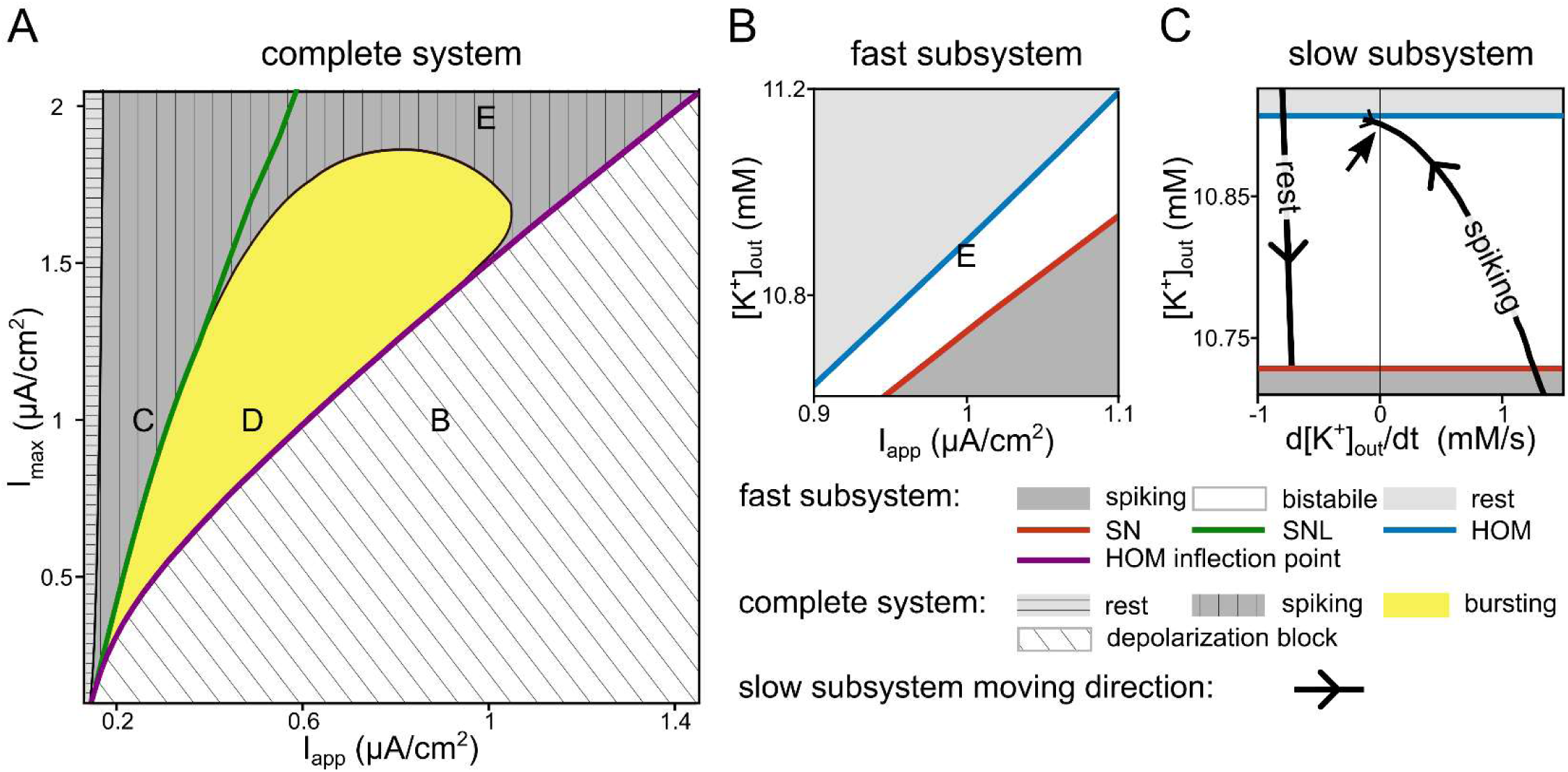
Dynamics of the complete system for varying pump density. Depending on the maximum ATPase pump current (I_max_) and applied current (I_app_), the complete system can show different dynamics (rest, tonic spiking, bursting, and depolarization block). This figure focuses on a low potassium concentration. (**A**) Bifurcations in the complete system as well as the corresponding fast subsystem saddle-node-loop (SNL) and HOM inflection point. The regions labelled B, C, and D in this figure correspond to the same parameter space as shown in Fig 6 with the same labelling. The area between the saddle-node-loop (SNL) and HOM inflection point lines of the fast subsystem represents the parameter space where bursting can occur according to the fast subsystem analysis. Only the yellow zone within this region displays bursting behaviour. Numerically, the boundary of the bursting region is detected as the transition of the inter-spike interval distribution from unimodal (tonic spiking) to bimodal (bursting). High pump densities lead to tonic spiking instead of bursting. (**B**) Two-parameter (extra cellular potassium, [K^+^]_out_, and I_app_) fast subsystem bifurcation diagram for the case indicated by E in panel A. In the fast subsystem of this case, the necessary condition for bursting, i.e., a bistable region bounded by homoclinic (HOM) and saddle node (SN) bifurcations from above and below, respectively, is satisfied. (**C**) Reduced slow subsystem dynamics overlaid onto the bifurcation map of fast subsystem corresponding to letter E in panels A and B. The spiking branch of the reduced slow subsystem intersects the line d[K^+^]_out_/dt=0, indicated by the black arrow. This intersection signifies the presence of a stable fixed point on the spiking branch within the bistable region of the fast subsystem. Consequently, the complete system exhibits tonic spiking, despite the fast subsystem meeting the conditions for bursting. I_max_=2 μA/cm^2^, I_app_=1 μA/cm^2^.

For I_app_ higher than the HOM inflection point (purple square in Fig 6A), the complete system dynamics ends up in depolarization block. After passing through a Hopf bifurcation, the neuron loses the ability to spike in a fold of limit cycles (FLC) bifurcation. This can be referred to as a depolarization block. The letter B in Fig 6A corresponds to one example that converges towards depolarization block. If the system starts in low [K^+^]_out_ at position B, i.e., the tonic spiking region of the fast subsystem (dark grey area), with each spike, [K^+^]_out_ increases, moving the system upwards along flow field direction of the complete system (filled black arrows in Fig 6A). The system passes the bistable zone but does not encounter the HOM bifurcation. The limit cycle terminates in an FLC bifurcation instead. The same can also be seen in the dynamics of the reduced slow subsystem for the same modelling parameters (Fig 6B). Here, if we start from the spiking branch (solid line), the time derivative of [K^+^]_out_ is positive. Hence, the system slides on the spiking branch from lower to higher [K^+^]_out_, until it reaches the FLC bifurcation. At this point, the spiking branch terminates, and the system jumps to the resting branch with a positive time derivative of [K^+^]_out_ instead of the normal resting state. The time derivative of [K^+^]_out_ on this branch is positive because this state corresponds to an upstate voltage at which more voltage-gated channels are in a conducting state. With the given I_max_, the pump is insufficient to compensate and [K^+^]_out_ continues to increase until the silent branch eventually approaches zero in a very large [K^+^]_out_. If the system starts at [K^+^]_out_ higher than the SN bifurcation of fast subsystem and is on the other resting branch of the slow subsystem (left side of the Fig 6B), however, the negative time derivative moves the system along this branch in the direction of decreasing [K^+^]_out_. As it progresses, the system reaches the SN bifurcation, transitioning into a spiking state, and eventually leading to the depolarization block.

Note that, in Fig 6A, for I_app_ lower than the HOM inflection point of the fast subsystem and [K^+^]_out_ higher than HOM bifurcation in the bistable region of the fast subsystem, the complete system dynamics are similar to the dynamics of Figure 6B, where [K^+^]_out_ is higher than the fast subsystem SN bifurcation. Therefore, two directions for the complete system dynamics exist. The first one increases [K^+^]_out_, resulting in depolarization block. The second one decreases [K^+^]_out_, in contrasts to the scenario in Fig 6B, resulting in either tonic spiking or bursting, in dependence of the size of I_app_. Hence, the dynamics of the complete system exhibits bistability (depolarization block versus either bursting or tonic spiking).

In this section, we demonstrated that our basic neuron model possesses the ability to not only generate slow rhythmic bursting activity but also to exhibit other significant dynamics such as tonic spiking and depolarization block. Each of these dynamics plays a crucial role in both physiological and pathological scenarios.

#### Dependence of the bursting mechanism and other dynamics on the pump density

Since the Na^+^/K^+^-ATPases are crucial elements in the feedback mechanism underlying the slow dynamics, the pump density (which in our model can be determined by I_max_) is a natural parameter to explore the robustness of the described mechanism (regarding both existence and extent of the bursting region). The complete system dynamics is next explored in the I_max_-I_app_-plane (Fig 7A).

In the complete system dynamics, four distinct regions are discernible: rest, tonic spiking, bursting, and depolarization block. The same description employed for the analysis of Fig 6A can be applied here as well. When I_app_ is below the HOM inflection point and [K^+^]_out_ above the HOM bifurcation in the bistable region of the fast subsystem, the complete system dynamics exhibits bistability between what is demonstrated in Fig 7A and depolarization block. For completeness, we note that chaotic behaviour is observed during simulations in the bursting region near the border where dynamics in the complete system change. The chaotic action potentials that is added at the end of the spiking phase can happen in fold/homoclinic bursters and has been studied before [29]. Regarding the impact of I_max_, the pump density, the following relations can be deduced from the figure: The larger I_max_, the larger the applied current needs to be to reach the depolarization block. Over a large range of I_max_, the ranges of tonic spiking as well as bursting expand with I_max_, before the bursting region shrinks and terminates. Note that in the previous sections, we examined representative cases from each region of depolarization block, tonic spiking, and bursting from the preceding sections (see Fig 3,4, and labels B, C, and D in Fig 6). B, C, and D share the same I_max_ value, which was also used for the two-parameter bifurcation diagram in Fig 6A.

The fast subsystem SNL bifurcation and the HOM inflection point are denoted by green and purple lines, respectively, in Fig 7A. In the preceding section, we established their pivotal role in shaping the slow burst dynamics. The bistable region of the slow subsystem, existing between these two lines, serves as a necessary condition for bursting dynamics, yet it does not suffice for the induction of bursting behaviour. Bursting in the complete system occurs solely within the yellow region bounded by the SNL bifurcation and the HOM inflection point lines. In the remaining area between these two lines, although the fast subsystem fulfils the requirements of bursting (bistability between tonic spiking and quiescence, and distortion due to the presence of the pump), the slow subsystem’s hysteresis loop fails to form, leading the entire system into tonic spiking instead of bursting. The maximal pump density (I_max_) thus defines a region in which the complete system bursts.

We conducted a case study at the position marked E in Fig 7A, where the fast subsystem is bistable, yet non-bursting (because the pump strength is too strong for a hysteresis loop to develop). The corresponding slow-fast analysis diagrams are shown in Fig 7B and C. Here, the fast subsystem exhibits a bistable area bounded between quiescence and tonic spiking in high and low [K^+^]_out_, respectively. While the fast subsystem fulfils essential prerequisites for bursting, the complete system does not travel along the bistable region of the fast subsystem. Instead, it tonically spikes (corresponding to position E in Fig 7B) because the stable fixed point of the reduced slow subsystem on the spiking branch destroys the hysteresis loop (Fig 7C). We can conclude that if the pump density is too high, bursting can be evaded and the system stabilizes in a tonic spiking state.

From this section, it becomes evident that the pump plays a vital role in shaping the diverse dynamics that are possible within a model with spike generating dynamics of type Ⅰ, when the dynamics of extracellular potassium are taken into account. While the pump can establish the necessary conditions for bursting, it can also suppress bursting incidents. Additionally, higher pump density contributes to postponing depolarization block.

Finally, we briefly note that the mechanism was also tested in the presence of dynamic Na^+^ concentrations. To this end, we modified the model by incorporating intracellular sodium concentration ([Na^+^]_in_) dynamics through Eqs 11 and 12 and by adjusting the ATPase equation to also account for changes in [Na^+^]_in_ (see Eq 13). The result is shown in S2 Fig. The desired bursting mechanism is present in the modified model, too. However, including [Na^+^]_in_ dynamics in the model can introduce more intricate behaviours. For example, in panel A of S2 Fig, sodium slightly accumulates over time and can result in alterations of spike amplitude and spiking threshold [30]. These dynamics, however, are not the focus of this paper.

## Discussion

In this article, we demonstrate that a neuron whose dynamics of the extracellular potassium are taken into account requires only two essential elements to exhibit slow bursting: a bistable region arising from an SNL bifurcation and a feedback loop mediated by a Na^+^/K^+^-ATPase. In contrast to previous models for slow bursting, this firing mode requires neither ion channels with slow dynamics nor rhythmic inputs. Our slow-fast bifurcation analysis reveals a ubiquitous mechanism: square-wave bursting via a fold-homoclinic hysteresis loop, which, in principle, can be encountered in all type I excitable neurons with spike onset via a SNIC bifurcation, the spiking dynamics of which have been shown be tuneable into an SNL regime via a number of physiological parameters [27,28]. The evidence presented assigns a pivotal role to the Na^+^/K^+^-ATPase pump in the generation of slow bursting as well as depolarization block. The exploration of the mechanistic core and robustness of the described bursting mechanism described here provides novel perspectives for the role of elevated extracellular potassium levels in neural pathologies such as seizures or spreading depolarization and potentially their treatment.

### Accumulation of extracellular potassium concentration serving as a feedback signal

Our study demonstrates that the interplay between potassium dynamics and neuronal excitability in class Ⅰ neuron models with SNIC dynamics can give rise to slow bursting. The general idea of generating diverse spiking dynamics via feedback from increased extracellular potassium is not new (for modelling studies see, for example, [12,23,31–37]) and supported by experimental evidence. For example, fluctuations in extracellular potassium levels have been observed to co-vary with EEG and FP waves during sleep or seizures [21,22,38]. Increases in extracellular potassium have been reported during pathological conditions like seizures and spreading depression [39–41]. Moreover, prior studies have shown that accumulated extracellular potassium does not rapidly diffuse away from neurons, thereby providing a feedback signal that can influence neuronal dynamics [42–46]. Previous modelling work has focused predominantly on mechanisms based on ion channels with dynamics slower or more complex than those of classical Hodgkin-Huxley Na and K channels [47], or, alternatively, concentration dynamics paired with the classical Hodgkin–Huxley channels were augmented by additional features like multiple compartments or homeostasis mechanisms. Our findings show, however, that none of these factors are strictly necessary requirements for rhythmic activity. In contrast, the minimal mechanism described here has not received sufficient attention.

### The relevance of the SNL bifurcation in voltage dynamics

At the core of the mechanism is the SNL bifurcation and the bistable region initiating from it in the fast subsystem. The latter is bounded by the HOM and SN bifurcations, two codimension-one bifurcations that appear from the codimension-two SNL bifurcation. Proximity of an SNL bifurcation can be found in a generic class of neuron models, i.e. all models starting with class I excitability [16,28], which encompass a significant proportion of neuron models, ranging from isolated gastropod somata to mammalian hippocampal neurons [27]. Note that in the bistable region between the HOM and SN bifurcations, the presence of noise in the system or its input can result in stochastic bursting that emerges from switching between the stable limit cycle attractor and the fixed-point dynamics (see, for example [30]). Here, in contrast, the bursting mechanism is deterministic and arises from the Na^+^/K^+^-ATPase-mediated feedback. Specifically, the activity of the ATPase results in a distortion of the bifurcation diagram, which is the crucial effect that enables the deterministic bursting. Deterministic bursting via this mechanism cannot occur in class II neurons because they lack the SNL bifurcation.

### Dynamical potassium concentration sets four dynamical regimes in one minimalistic neuron model

The minimal biophysical model also demonstrates the ability to exhibit both bursting and depolarization block, which aligns with previous observations indicating that seizures and spreading depolarizations (SD) can originate from the same neuronal population [34,37]. Depending on Na^+^/K^+^-ATPase density and input strength, the complete concentration-dependent system exhibits four different regimes: rest, tonic spiking, bursting, and depolarisation block, which have been associated with a range of healthy and/or pathological states. For example, seizures can be triggered by a handful of neurons going through bursts of activity [48]. In SD, the phase of neuronal hyperactivity preceding the depression can arise from synchronized activity of bursting neurons, as observed by population spikes in electrocorticography (ECoG) traces [49]. In the wave of death (i.e. the massive and simultaneous depolarization of neurons [50]) or when the SD wave changes from hyperactivity to silence, depolarization-block-like dynamics could be observed [34]. The changes in extracellular potassium concentration during bursting and depolarization block in our model are consistent with experimental observations during seizures and SD, as well as during sleep rhythms [21,22,51].

### The relevance of the Na^+^/K^+^-ATPase

Our findings assign an important role to Na^+^/K^+^-ATPases [52] in shaping a neuron’s dynamics. The pump and its isoforms are known as a generic player in the homeostasis of extracellular potassium, resting potential, as well as cell volume. Experimental evidence indicates that neuronal Na^+^/K^+^-ATPase function extends beyond the stabilization of extracellular potassium. Along these lines, it has been argued that if most of the potassium secreted by neurons were siphoned off by astrocytes, neuronal activity could not continue for tens of seconds because of imbalances in potassium concentration [53]. Our model indicates yet another function of the Na^+^/K^+^-ATPase: as an enabler of rhythmic activity when extracellular potassium concentration is elevated. Previously, studies have introduced the role of Na^+^/K^+^-ATPase in rhythmic bursting activity in pattern generators both experimentally and theoretically [54,55]. However, these studies primarily emphasized the interaction between h-current and the ATPase pump in controlling Na^+^ and did not investigate the burst mechanism proposed in this paper. The involvement of Na^+^/K^+^-ATPase in the induction of rhythmic activity is consistent with previous proposal, which indicates that any [K^+^]_out_ or voltage-dependent ion channels that elicit inward currents can generate seizure-like rhythmic activity, provided that the inward current (secondarily) triggers the release of K^+^ into a confined extracellular space [31] – a set of conditions met by Na^+^/K^+^-ATPase.

Our model prediction that a higher pump density shifts the initiation of the bursting mode and depolarization block towards higher input currents agrees with the experimental observation that the Na^+^/K^+^-ATPase helps to inhibit seizures and SD. Specifically, inactivation of the neuron-specific alpha3-isoform of Na^+^/K^+^-ATPase has been shown to induce seizures in mice [56]. Further, the loss-of-function mutation that causes chorea-acanthocytosis is accompanied by impaired capacity of the Na^+^/K^+^-ATPase in neurons, and epilepsy is among the symptoms of this neurodegenerative disorder [57]. Even partial inactivation of the Na^+^/K^+^-ATPase with ouabain has been shown to cause SD-like depolarization: An epileptic population spike is followed by an ouabain-induced SD [58], which is consistent with the cellular dynamics observed in our model. Moreover, a shortage of oxygen or ATPase, as it may, for example, occur during ischemia, renders the pump less effective. We demonstrate that an insufficient pump current can first induce bursting and then facilitates depolarization block. This agrees with depolarization events occurring in hypoxic and ischemic conditions [58,59] and previous suggestions that hypoxia or weakened pump current can drive neurons into seizure states [31,60]. In addition, an experimental study has revealed a correlation between seizures and specific oxygen pressure levels, inferring that a greater reduction in oxygen pressure disrupts ATPase pump function, resulting in the cessation of seizure-like activities [61]. Our model, in alignment with the experimental findings, indicates that – although an inadequate pump current can trigger seizures (bursting) – a reduction in pump activity can also contract the bursting (and thus seizure-relevant) region within the parameter space.

### The relevance of potassium homeostasis

The dynamics described in this study do not explicitly consider other ionic transporters, or external potassium regulators, such as diffusion, or glial cells [62–66]. The model’s extracellular space can, however, be interpreted as an effective compartment that also subsumes possible additional potassium homeostasis mechanisms. Hence, the pathological dynamics introduced in our mechanism could arise from anomalies in these kinds of regulators. This hypothesis is consistent with the experimental observation that dysfunctional astrocytes are crucial players in epilepsy, assuming that extracellular potassium homeostasis is impaired as a function of the astrocytic impairment [67]. Other experiments have implicated dysfunctional astrocytes in SD [68]. We note that although our model produces intriguing dynamics at the level of a single cell, it cannot generate some of the more complex behaviours, such as recovery from depolarization block. This recovery requires the explicit involvement of external potassium regulators [33], exceeding the scope of this paper.

## Summary

In summary, we present a minimal mechanism for deterministic bursting activity that arises from the interplay of very common spiking dynamics with changes in the extracellular potassium concentration mediated by a ubiquitous Na^+^/K^+^-ATPase. While other, more complex models, exhibit deterministic bursting too, the mechanism described in this study relies only on a minimal set of physiological assumptions in neurons and its dynamics agree with experimental observations in pathological states like epilepsy or spreading depolarization. The underlying mechanism strengthens the role of Na^+^/K^+^-ATPases in the generation of slow rhythms and their relevance as a therapeutic target in pathology. While more complex models may capture bursting dynamics faithfully, the minimal model enabled us to deduce the exact mechanism of the bursting regime.

We anticipate that this detailed understanding, as well as the efficiency of the model in numerical simulations, will facilitate analyses aimed at mechanistically linking the biophysics of neurons to the behaviour of the networks that embed them.

## Methods

### Model description

In the following, we describe the parsimonious biophysical model, composed only of two fast spike generating currents and a Na^+^/K^+^-ATPases pump as well as extracellular potassium concentration ([K^+^]_out_) dynamics.

Our model is based on the Wang-Buzsáki model [69]. As it is shown in Fig 1, the model consists of two compartments: the neuron and its’ extracellular space. The outward potassium current through the neuron’s voltage-dependent channels will increase the concentration of [K^+^]_out_. Conversely, [K^+^]_out_ is decreased by the Na^+^/K^+^-ATPase pump activity. We calculated these changes by the mass conservation equation of [K^+^]_out_. Variation in [K^+^]_out_ affects the reversal potential of the potassium, which is calculated by the Nernst equation.

#### The Wang-Buzsáki model

The Wang-Buzsáki (WB) model is a modified version of the famous Hodgkin-Huxley (HH) model. However contrary to the HH model for invertebrate motoneurons, the WB model is categorized as a class Ⅰ neuron model, mimicking fast-spiking vertebrate interneurons dynamics. The model has two fast spike-generating sodium and potassium voltage-dependent channels [69]. The potassium and sodium currents are described below:

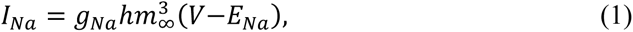

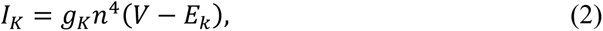

Where sodium channel inactivation variable h and potassium channel activation variable n are defined with the following equations:

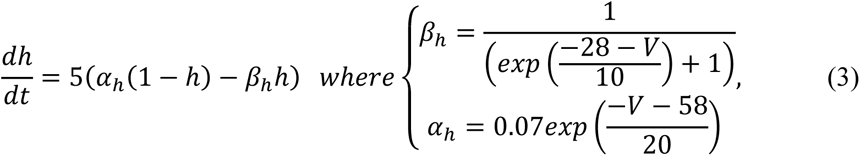

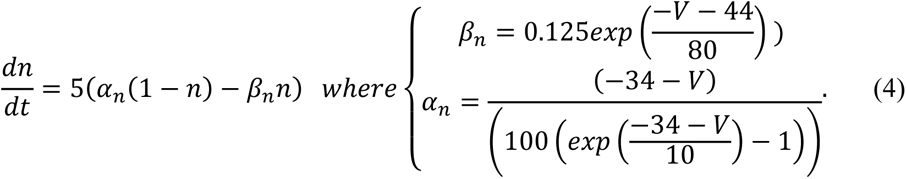

Within the framework of the Wang-Buzsáki model, the activation variable of the transient sodium current (m) is considered to evolve rapidly and is thus replaced with its corresponding steady-state function. This approach serves as a means to simplify the model, transitioning it from a four-dimensional system to a three-dimensional one.

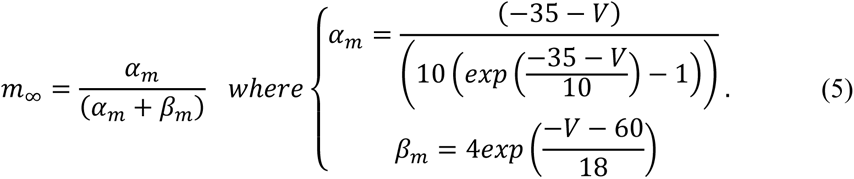

The Wang-Buzsáki model also has a leak current, which is described below:

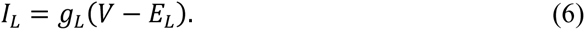

All of the symbols and constant of this section is described in the Table 1.

**Table 1.** Nomenclature.

| *Symbol* | *Names* | *Values (unit)* |
| --- | --- | --- |
| $[K^+]_{out}$ | extracellular potassium concentration | variable (mM) |
| $\alpha$ and $\beta$ | gating variables | variable ($ms^{-1}$) |
| $A_{cell}$ | surface area of cell | $3.142 \times 10^{-6}$ ($cm^2$) [70] |
| $C$ | membrane capacitance | 1 $\mu F/cm^2$ [69] |
| $E_k$ | potassium reversal potential | variable (mV) |
| $E_L$ | leak reversal potential | -65 (mV) [69] |
| $E_{Na}$ | sodium reversal potential | 55 mV [69] |
| $F$ | Faraday's constant | $9.694 \times 10^4$ (C/mol) [70] |
| $g_k$ | potassium conductance | 9 (mV) [69] |
| $g_L$ | leak conductance | 0.1 ($mS/cm^2$) [69] |
| $g_{Na}$ | sodium conductance | 35 ($mS/cm^2$) [69] |
| $h$ | sodium channel inactivation | variable ____ |
| $I_K$ | potassium current | variable ($\mu A/cm^2$) |
| $I_L$ | leak current | variable ($\mu A/cm^2$) |
| $I_P$ | $Na^+/K^+$-ATPase pump current current | variable ($\mu A/cm^2$) |
| $I_{app}$ | Injected (stimulation) current | varies per simulation ($\mu A/cm^2$) |
| $I_{max}$ | $Na^+/K^+$-ATPase pump maximal current | varies per simulation ($\mu A/cm^2$) |
| $I_{Na}$ | sodium current | variable ($\mu A/cm^2$) |
| $m_\infty$ | fast sodium channel activation | variable ____ |
| $n$ | potassium channel activation | variable ____ |
| $r_v$ | ratio of extracellular to intracellular volumes | 0.15 ____ [70] |
| $t$ | time | variable (s) |
| $V$ | membrane potential | variable (mV) |
| $V_{cell}$ | extracellular volume | $5.23 \times 10^{-10}$ ($cm^3$) [70] |

#### Na^+^/K^+^-ATPase pump current

To generate action potentials in neurons, the presence of ionic gradients between intracellular and extracellular space is required. This entails a high concentration of K^+^ within the cell and a low concentration outside, while Na^+^ maintains a low concentration within the cell and a high concentration outside. To maintain the resting potential and neuronal excitability the cell membrane contains pumps like the Na^+^/K^+^-ATPase. The Na^+^/K^+^-ATPase pump puts 3 sodium ions out of the cells while pumping 2 potassium ions into the cells; hence, this pump is not electroneutral. The concentration dependence of the pump current is an sigmoidal function, adapted from Hübel et al. [33] and modified to match our model so the resting potential of the Wang-Buzsáki model is not changing, dramatically.

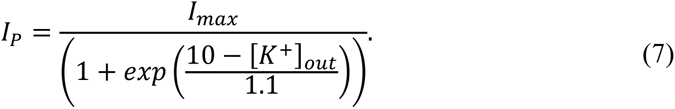

This pump equation demonstrates the adaptability of the pump current in response to changes in extracellular potassium concentration. The more potassium has accumulated outside of the cell, the stronger the pump current that pumps potassium back inside. Nevertheless, the pump’s current never exceeds the maximum level, I_max_ set by the density of pumps in the membrane.

#### Membrane potential equation

Having described all the currents of our model, we can now calculate membrane potential dynamics as below:

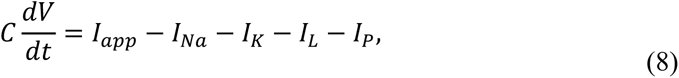

Where I_app_ is the applied current or stimulus.

#### Ionic continuity equation

Neuronal activity raises the potassium reversal potential, while lowering the sodium reversal potential. In general sodium reversal potential have a strong impact on action potential height, while the potassium reversal potential induces bifurcations of the resting state and the saddle that determines the threshold manifold. In particular homoclinic connections to the saddle are affected by changes in potassium reversal potential [30]. Furthermore, due to higher potassium conductance than sodium conductance at the resting potential, the resting membrane potential is typically closer to the potassium reversal potential. Additionally, due to the smaller extracellular space in relation to intracellular space, changes in extracellular potassium concentration ([K^+^]_out_) can have a greater effect on neuronal behaviour than any other neuronal concentration[23]. Therefore, it is reasonable to focus on extracellular potassium dynamics in our model.

For converting the potassium current to potassium ionic flux, we multiply the pump current by two; as in each pumping cycle, two potassium cations are exchanged. Thus, the continuity equation of [K^+^]_out_ is as follows:

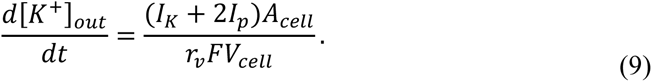

In our model, the reversal potential of potassium (in Eqs 2) is dynamic and it is calculated according to the Nernst equation [33,70].

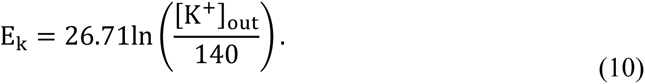

### Dynamic intracellular sodium

To demonstrate that the slow burst dynamics observed in our model are persistent, we have added sodium concentration dynamics to the model. The results of this addition are reported in the Supplement materials (S2 Fig).

The extracellular sodium concentration is higher than the intracellular sodium concentration ([Na^+^]_in_). Additionally, the Na^+^/K^+^-ATPase pump is responsive to changes in [Na^+^]_in_. Hence, to enhance our model with sodium dynamics, we can focus on integrating the dynamics of [Na^+^]_in_.

For simulating the model with dynamics [Na^+^]_in_, we add the continuity equation for [Na^+^]_in_ to the model:

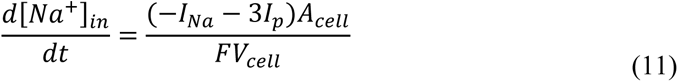

We also calculate the reversal potential of sodium with the equation below instead of using it as a parameter.

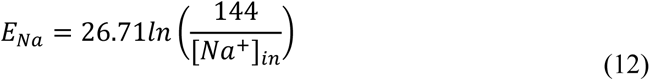

Finally, we use the complete model of the pump [33] with some modifications.

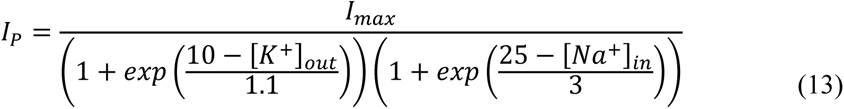

To reproduce the same bursting pattern as shown in Fig 2, we need to set [Na^+^]_in_ to approximately 18.4 mM. Using this concentration in Eq 12 results in E_Na_=55 mV, which aligns with our previous simulations. Moreover, in 13, the I_max_ should be adjusted. With [Na^+^]_in_ near 18.4 mM, I_max_ should be around 10 µA/cm^2^. This adjustment allows us to conduct the model simulation under nearly the same conditions that closely resemble the results shown in Fig 2.

The outcome of the simulation incorporating all the modifications above is presented in panel A of S2 Fig. With the [Na^+^]_in_ oscillation around 18.4 mM, we successfully replicate the nearly identical bursting dynamics showcased in Fig 2. However, it’s worth noting the presence of a gradual drift in the sodium concentration.

For another sanity check, in the modified model with dynamic [Na^+^]_in_, we set [Na^+^]_in_ =18.4 mM. The outcome of this simulation is depicted in panel B of S2 Fig, showcasing identical results to those seen in Fig 2. Hence, the burst dynamics described in this paper can manifest in more complicated systems.

### Slow-fast method

The deterministic bursting introduced in this paper results from the interaction between dynamics on different timescales. First, the fast spiking timescale mostly determined by gating variables of channels and the membrane time constant (Eqs 3, 4, and 8). Second, the slower timescale of the concentration dynamics. The slow dynamics is represented by variation in [K^+^]_out_ (Eq 9).

To gain a better understanding of our system’s behaviour, we analysed our neuron model using the method of timescale separation [10,24]. For this one need to assume that slow and fast timescales are sufficiently separated. As a result, the fast subsystem perceives the slow subsystem ([K^+^]_out_) dynamics as a constant parameter. On the other hand, the oscillations of the spiking dynamics are so fast for [K^+^]_out_ dynamics that it only senses the average changes in the fast subsystem. By using this strategy, we can consider the fast and slow subsystems separately.

For the fast subsystem, we treat [K^+^]_out_ in the system of equations including Eqs 3, 4, and 8, as a constant variable and a bifurcation parameter. Then with the help of numerical continuation [71], we produce the bifurcation diagram of the fast subsystem (This is how we generated Fig 3, 5, 6A, and 7B).

To analyse the slow subsystem, we apply a numerical averaging method to obtain the reduced system [10,72]. The time evolution of slow subsystem ([K^+^]_out_) is given by Eq 9. But actually, the slow subsystem does not respond to the fast changes in the fast subsystem during the spiking phase but only perceives the average effect of each action potential. Therefore, for each desirable [K^+^]_out_, we can average the terms that relate to the fast subsystem (I_P_ and I_K_ in Eq 9) over one full action potential limit cycle, as shown below:

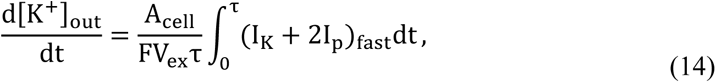

Where τ is the period of the fast subsystem’s limit cycle and the subscript “fast” shows that I_P_ and I_K_ belong to the fast subsystem in specific [K^+^]_out_.

For each spiking branch of the fast subsystem, we discretize the [K^+^]_out_ range that the branch covers and compute the [K^+^]_out_ derivative with averaging method (Eq 14) for each [K^+^]_out_ value. This allows us to calculate the reduced system for spiking branch of the fast subsystem.

When the fast subsystem is quiescent and on its’ stable fixed point, to determine the slow subsystem dynamics, it is enough that we take at the desired [K^+^]_out_, the fast subsystem’s fixed-point value of I_P_ and I_K_ and put it in Eq 9.

The result of the averaging method as reduced slow subsystem time derivative diagrams are shown in 4, 6B, 6C, and 7C.

The averaging method usually breaks down if the time scale difference between the slow and fast subsystems becomes too small. This situation can arise near the fast subsystem bifurcations, such as in the vicinity of the HOM bifurcation, where the spiking frequency becomes significantly reduced [24]. Hence, in these situations, we validate our results with numerical simulations to make sure the accuracy of our results.

### Numerical implementation

We performed numerical simulations using Python 3 [73] and the Brian 2 package [74]. The Rung Kunta fourth-order method was utilized with time steps of 0.005 ms, and in certain instances, adaptive time steps were employed. In case of fixed time step method, we verified the results with smaller time steps, which yielded consistent outcomes. Additionally, we carried out the continuation of the bifurcation diagrams using XPPAut [71] and Auto-07p [75] .

## Acknowledgments

We thank Robert Gowers, Louisiane Lemaire, and Philipp Norton for valuable feedback on the manuscript.

## Supporting Information

**S 1 Fig.**
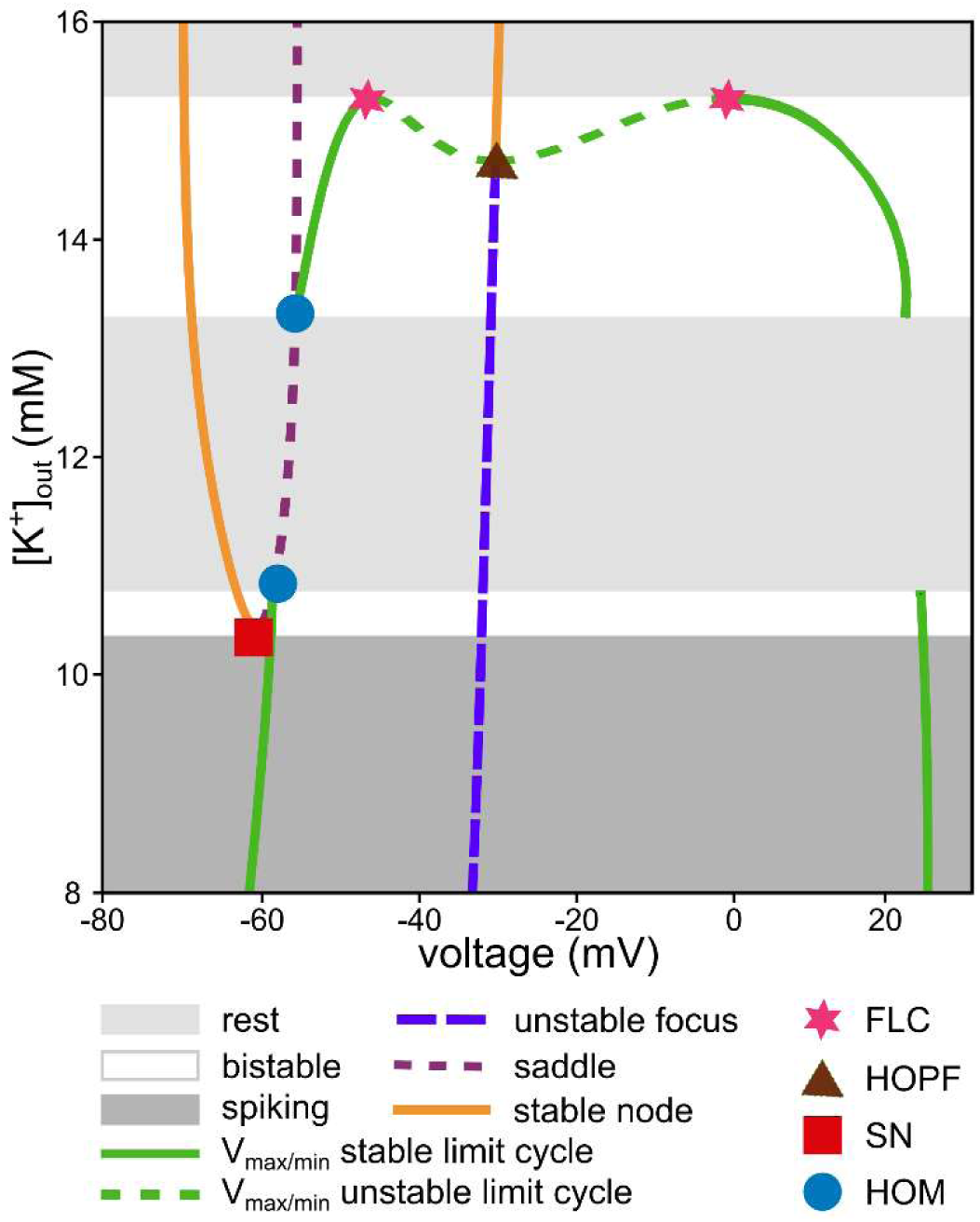
Fast subsystem bifurcation diagram around the bursting region. A zoom out of Fig 3B is depicted here. The bistable region between the saddle-node (SN) and homoclinic (HOM) bifurcation is the bursting region. There is a bistable region between HOM and Hopf bifurcations where a stable node and a stable limit cycle coexist. According to the complete system dynamics, the extracellular potassium concentration ([K^+^]_out_) is increasing if the fast subsystem spikes. On the stable branch of fast subsystem emerging from SN bifurcation, [K^+^]_out_ decreases. From the Hopf bifurcation of fast subsystem, another stable node is added to the fast subsystem. If the fast subsystem is on this stable node, the complete system dynamics dictates an increase of the [K^+^]_out_.

**S 2 Fig.**
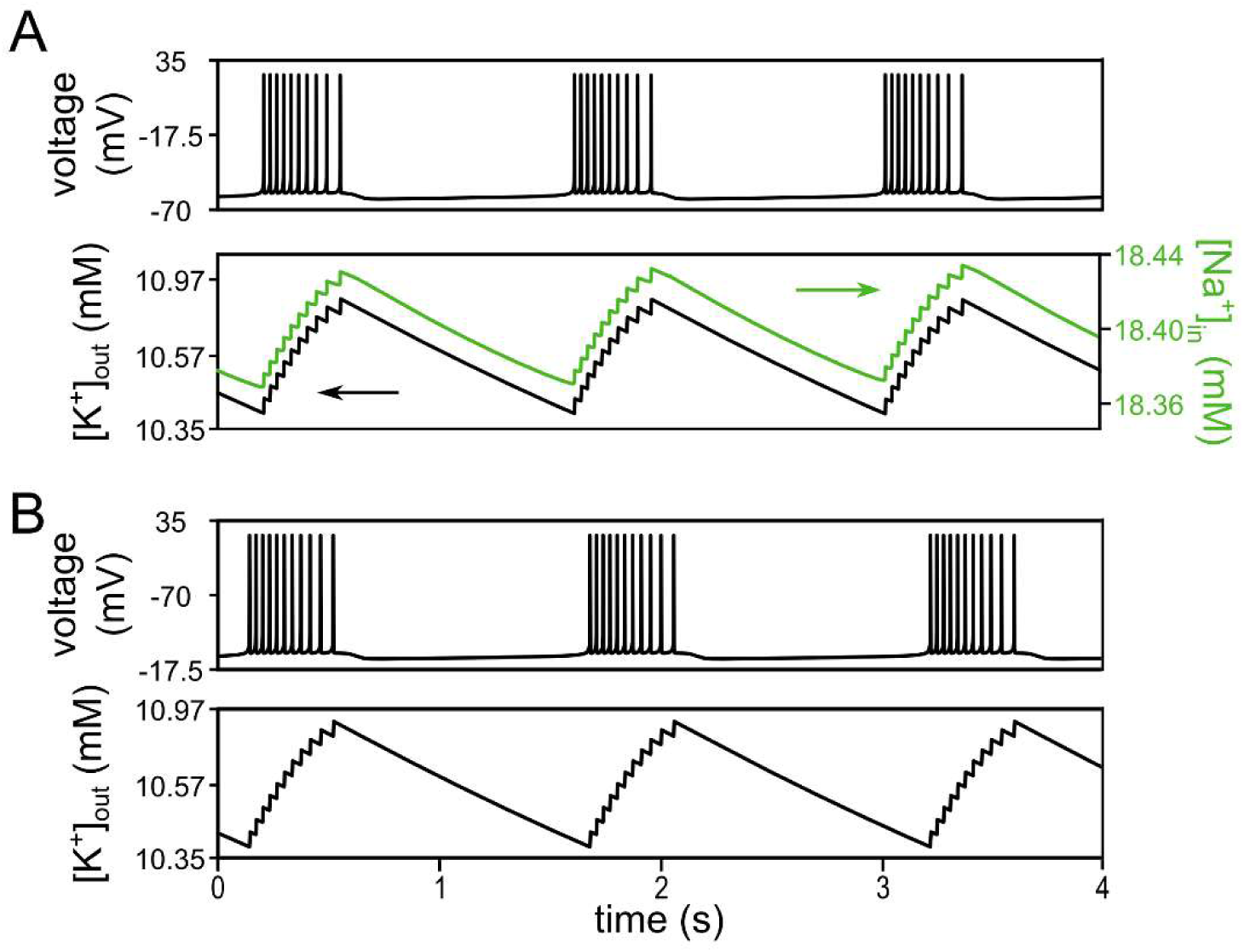
Effect of intracellular sodium on bursting dynamic. The bursting mechanism introduced in this paper is not affected by the dependence of the ATPase pump on [Na^+^]_in_. (**A**) Voltage and ionic concentration dynamics during bursting in a modified model with dynamic [Na^+^]_in_. Despite the addition of [Na^+^]_in_ dynamics, the bursting described in this paper is still observed, with extracellular potassium accumulating during spiking and decreasing during rest, similar to the behaviour shown in Fig 2. (I_app_=0.5 μA/cm^2^, I_max_=10 μA/cm^2^.) (**B**) Bursting dynamics for the modified model with fixed [Na^+^]_in_. By fixing [Na^+^]_in_ at 18.4 mM (mean approximation from A) in the modified model, the bursting dynamic is the same as what is depicted in the main model of this paper (see Fig 2).

## Data availability statement

All relevant data are within the paper, its Supporting Information files, and on Zenodo at (DOI will be provided upon acceptance).

